# Sustained Improvements in Student Outcomes Following Integration of Clinical Case Narratives in Veterinary Microbiology Curriculum

**DOI:** 10.64898/2026.05.22.727235

**Authors:** Aria Eshraghi, Lauren K Logsdon

## Abstract

Microbiology education in veterinary curricula requires students to integrate complex foundational knowledge with clinical application, yet traditional lecture-based approaches often emphasize memorization over higher-order reasoning. In this study, we evaluated the impact of integrating clinically oriented, case-based instruction into a veterinary microbiology course within a Doctor of Veterinary Medicine curriculum. Using a quasi-experimental, multi-year design, student outcomes were compared before (2019, 2021) and after (2022–2025) implementation of case-based teaching while maintaining consistent course content, structure, and assessments. Introduction of clinical case examples was associated with significant and sustained improvements in student evaluations across multiple domains, including perceived relevance, critical thinking, and overall course value. Instructor-related evaluation metrics also improved. Student performance, measured by final course grades, increased following the intervention without evidence of grade inflation. These findings demonstrate that integrating clinically relevant case narratives into microbiology instruction enhances student engagement and student performance. This work highlights a practical and scalable strategy for improving microbiology education, particularly within veterinary and other health-professions curricula.

## Introduction

Microbiology education in professional health curricula requires students to integrate large volumes of complex, detail-oriented information while simultaneously developing the ability to apply this knowledge in clinically relevant contexts. This challenge is particularly pronounced in veterinary microbiology, where students are required to understand a broad diversity of bacterial, fungal, and viral pathogens across multiple animal species, including their zoonotic implications within a One Health framework. The breadth and complexity of this material can increase cognitive load and may limit student engagement and deeper conceptual understanding when delivered through traditional lecture-based approaches^1^.

Conventional instruction in microbiology courses often emphasizes memorization of organism characteristics and disease associations. While this approach supports content coverage, it may not adequately promote higher-order cognitive skills such as application, integration, and clinical reasoning.^2–5^ Educational research has demonstrated that active learning strategies, including case-based instruction, can improve student engagement and facilitate the application of knowledge to realistic scenarios.^6–12^ By requiring learners to interpret complex information, distinguish relevant from extraneous details, and apply foundational concepts, case-based approaches may be particularly well suited for content-dense courses in professional programs.^4,7,13–15^

While many studies have demonstrated improved student outcomes and perception of learning using case-based approaches in science and medicine^11,13^, there remains limited longitudinal evidence evaluating the sustained impact of case-based instruction on both student perceptions and performance in veterinary microbiology curricula^7^. Previous studies on case-based learning in the field of Veterinary medicine employed case studies within a laboratory setting^16–18^ or in small group discussion sessions^5,19–22^. Few studies have examined the impact of case-based narratives incorporated into a large lecture format class and whether improvements in student evaluations are accompanied by measurable changes in student learning outcomes over multiple course offerings^23^.

In 2021, a second-year veterinary microbiology course within a Doctor of Veterinary Medicine curriculum was delivered using a traditional lecture format established by a previous instructor. In 2022, the course was redesigned to incorporate clinically oriented case narratives throughout lectures. These cases were developed or adapted from clinical reports and were intentionally designed to be complex and include extraneous information, requiring students to engage in higher-order reasoning and apply microbiological principles in diagnostic and therapeutic contexts. Importantly, course content, learning objectives, and overall structure remained consistent across years, allowing comparison of student outcomes before and after the instructional change.

The objective of this study was to evaluate the impact of incorporating clinical case-based instruction on student engagement and performance in a veterinary microbiology course. Specifically, we asked: (i) Does implementation of case-based teaching improve student evaluations of instructor effectiveness and course value? and (ii) Is this change associated with improvements in student performance, as measured by course grades? We hypothesized that introduction of clinically relevant case examples would result in immediate and sustained improvements in student evaluations and would be associated with increased student performance without evidence of grade inflation.

## Methods

### Course context and instructional design

This study was conducted in a veterinary microbiology course within a Doctor of Veterinary Medicine (DVM) curriculum at University of Florida. The course covers fundamental aspects of bacteriology, mycology, and virology, with emphasis on pathogen characteristics, disease mechanisms, laboratory diagnosis, antimicrobial therapy, and clinical relevance within a One Health framework.

From 2019 through 2021, the course was delivered using a traditional lecture-based format established by a previous instructor. Pathogens were presented in groups based on shared characteristics, and each organism was taught using a standardized structure that included nomenclature, physical characteristics, virulence factors, associated diseases, diagnostic approaches, and relevant information on vaccines and treatment. In 2021, a new course coordinator was appointed, and the five-member teaching team delivered the course using these existing materials without substantive modification. Data from 2020 were excluded from analysis due to substantial changes in course delivery associated with the COVID-19 pandemic, during which instruction was conducted primarily online. Student assessment structure was consistent across all years and included ten quizzes and four examinations distributed throughout the course. These assessments were designed to evaluate both foundational knowledge and critical thinking skills through the application of microbiological concepts. Assessments remained consistent throughout the study period, with only minor modifications for clarity or formatting.

### Instructional intervention

Beginning in 2022, a significant instructional intervention was implemented without altering course content, learning objectives, or overall structure. Instead, the method of content delivery was modified to incorporate clinically oriented case examples throughout lectures. These case examples were developed by the course instructors or adapted from published case reports and were intentionally designed to provide vivid and complex examples that were reflective of real-world clinical scenarios. Cases frequently included extraneous or distracting information to simulate the ambiguity and complexity encountered in veterinary practice. Students were required to identify relevant clinical details, integrate microbiological knowledge, and apply concepts to diagnostic and therapeutic decision-making.

Case examples were delivered using a progressive disclosure format designed to simulate clinical decision-making. Students were initially presented with partial clinical information, including patient history and environmental context, which intentionally incorporated both relevant and extraneous details reflective of real-world veterinary practice. At this stage, students were prompted to generate differential diagnoses and consider whether the etiology was infectious, non-infectious, or multifactorial. Additional information, including physical examination findings and epidemiological patterns (e.g., involvement of multiple animals, lesion progression), was then introduced incrementally to refine diagnostic reasoning. Only after discussion of these elements was the causative organism revealed, followed by targeted instruction on microbial virulence factors, diagnostic approaches, transmission, and control strategies. An example of this instructional approach is provided in Supplemental figure S1.

The course was co-taught by four to five faculty members across all years of the study period. Three faculty members delivered content in bacteriology and mycology, while one to two faculty members taught virology. Despite the multi-instructor format, the incorporation of case-based examples was implemented across the course beginning in 2022. Importantly, the number of assessments, grading structure, and overall course organization remained unchanged, allowing differences in student outcomes to be attributed primarily to changes in instructional delivery.

### Data collection

Student evaluation data were collected at the conclusion of each course offering prior to the release of final grades. Evaluations were administered using a standardized institutional instrument with responses recorded on a 5-point Likert scale (1 = disagree, 5 = agree). Because the course was co-taught by multiple faculty members, reported Likert scores represent composite values averaged across all instructors contributing to the course in a given year. Ten evaluation items were analyzed in this study.

Six items assessed instructor performance:

Q3: The instructor was enthusiastic about the course.

Q4: The instructor explained material clearly and in a way that enhanced my understanding.

Q5: The instructor maintained clear standards for response and availability.

Q6: The instructor fostered a positive learning environment that engaged students.

Q7: The instructor provided prompt and meaningful feedback.

Q8: The instructor was instrumental to my learning.

Four items assessed course performance:

Q9: Course content was relevant and useful.

Q10: The course fostered interaction between student and instructor.

Q11: Course activities improved critical thinking.

Q12: Overall, this course was a valuable educational experience.

Student performance data were obtained in the form of final course grades for each year included in the study. Grades were analyzed as continuous variables.

### Study design and analysis

This study employed a quasi-experimental, pre-post design comparing student outcomes before (2019 and 2021) and after (2022-2025) implementation of case-based instruction. The year 2021 served as the primary baseline, as it reflects delivery of the course by the current coordinator and teaching team prior to the intervention. Composite scores were calculated for instructor-related items (questions 3-8) and course-related items (questions 9-12). Evaluation scores were summarized as mean and standard deviation values for each question and year. Departmental and college-level mean evaluation scores are presented for comparison; however, corresponding measures of variability (e.g., standard deviations) were not available to the authors and therefore are not reported.

Changes in evaluation scores and grades across years were assessed descriptively and through inferential statistical analyses. Comparisons between years were performed using Welch’s t-tests to account for unequal sample sizes and variances. Statistical analysis and graphing were performed using GraphPad Prism (Version 10.4.1)

This study was determined to meet criteria for a quality improvement project focused on instructional practices and did not involve identifiable human subjects’ data. As such, institutional review board approval was not required in accordance with University of Florida guidelines

## Results

### Course timeline demonstrates that instructional redesign, rather than changes in content or instructors, coincided with improved outcomes

Implementation of clinical case-based instruction in 2022 was associated with a marked and sustained improvement in both student evaluation scores and course performance. Core course content and assessments remained unchanged across years, allowing comparisons to focus primarily on differences in instructional delivery rather than changes in curriculum scope or grading standards. Course coordination transitioned from Coordinator #1 (2019– 2020) to Coordinator #2 beginning in 2021 (Figure 1).

**Figure 1.**
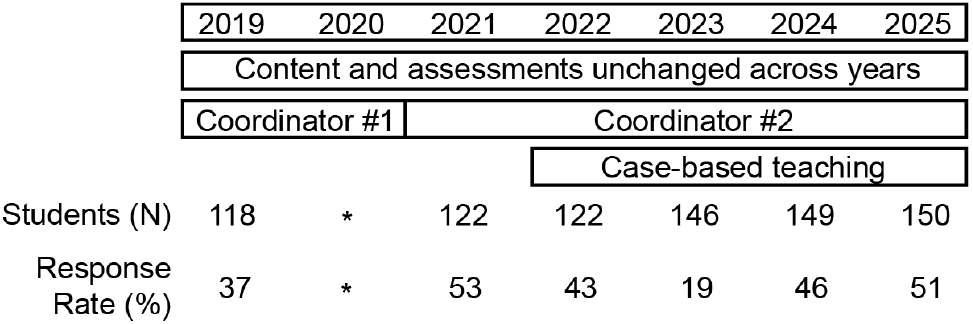
Course timeline and study design. Timeline of course delivery, instructional intervention, and enrollment across study years. The course was taught using a traditional lecture-based format from 2019-2021, with 2021 representing the baseline year under the current course coordinator. Clinical case-based instruction was introduced in 2022 and maintained through 2025. Core course content, structure, and assessments remained unchanged across years. Enrollment (N) and student evaluation response rates (%) are shown for each year. Data from 2020 were excluded from analysis due to COVID-19-related modifications to course delivery. Transition in course coordination is indicated.

Because the 2021 course was taught using the same general structure and materials as prior years, it served as the pre-intervention baseline. The major curricular change occurred in 2022, when case-based teaching was introduced and then maintained through 2025. This redesign emphasized clinically relevant scenarios while preserving the underlying microbiology content. The temporal association between this intervention and subsequent gains in student evaluations and course grades supports the hypothesis that presentation style, rather than content changes, contributed to improved outcomes.

Enrollment remained high throughout the study period, increasing from 118 students in 2019 to 150 students in 2025, indicating that improvements were achieved despite larger class sizes. Student evaluation response rates were variable but adequate in most years, ranging from 43% to 53% except for 2023 (19%). Data from 2020 were excluded because COVID-modified course delivery introduced substantial confounding variables.

### Course-related evaluation metrics improved significantly following implementation of case-based instruction

Beginning in 2022, traditional pathogen-focused lectures were supplemented with case-based instruction that embedded microbiological principles within realistic veterinary clinical scenarios. These cases were designed to integrate foundational microbiological concepts with realistic clinical scenarios, often including extraneous or distracting information (Supplemental figure S1). This approach required students to apply knowledge, prioritize relevant details, and engage in clinical reasoning, thereby shifting the focus from memorization to application. Together, these observations provide a foundation for examining how the intervention influenced specific domains of student evaluation.

Course-related evaluation scores demonstrated significant improvements following the 2022 intervention. Responses to question 9 (“Course content was relevant & useful”) increased significantly when comparing 2021 to all subsequent years, bringing scores in line with or exceeding college-level averages (Figure 2A). Notably, scores for this metric were similar between 2019 and 2021, suggesting that the improvement observed after 2022 reflects the change in instructional approach rather than pre-existing trends. A similar pattern was observed for questions 10 through 12, which assess interaction, critical thinking, and overall course value (Figures 2B-2D). Each of these measures showed statistically significant increases when comparing 2021 to 2022 and subsequent years, with sustained high performance through 2025. These increases elevated course evaluations to levels comparable to or exceeding departmental and college means. Collectively, these findings indicate that incorporation of clinical case examples was associated with broad improvements in student perceptions of course quality and educational value.

**Figure 2.**
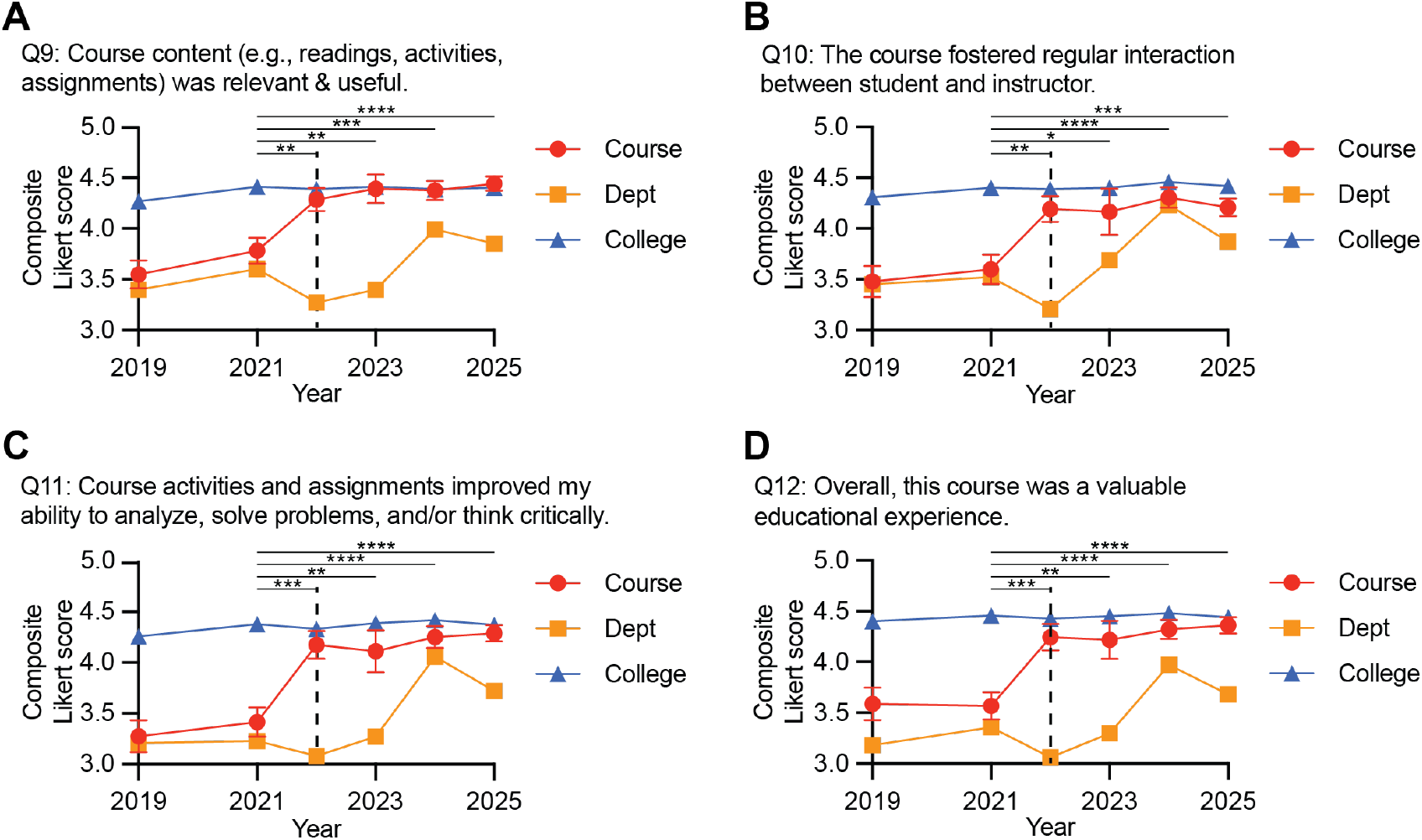
Course-related evaluation scores improved following implementation of case-based instruction. Composite Likert scores (mean ± SD, 1-5) for course-related evaluation items are shown across years for (A) relevance and usefulness of course content (Q9), (B) student-instructor interaction (Q10), (C) development of critical thinking (Q11), and (D) overall course value (Q12). Scores represent averages across all instructors for each course offering. Course-level scores (mean and standard deviation) are compared with departmental and college averages. Case-based instruction was introduced in 2022 (dashed line). Statistical significance was determined using Welch’s t-tests comparing each year to 2021 baseline (*p < 0.05, **p < 0.01, ***p < 0.001, ****p < 0.0001).

### Effect size analysis demonstrates moderate to large improvements in course-related outcomes

To quantify the magnitude of these changes, effect sizes were calculated using Hedges’ g with 2021 as the baseline. In general, Hedges’ g values of ≤0.2 are considered small or negligible, 0.2-0.49 small, 0.5-0.79 moderate, and ≥0.8 large. For question 9, comparison of 2021 to 2019 yielded a small and negative effect size, indicating no meaningful difference prior to the intervention (Figure 3A). In contrast, comparisons of 2021 to 2022 and subsequent years produced effect sizes exceeding 0.5, indicating moderate improvements. These values likely underestimate the true effect, as average Likert scale responses approached the upper bound of the scale in later years.

**Figure 3.**
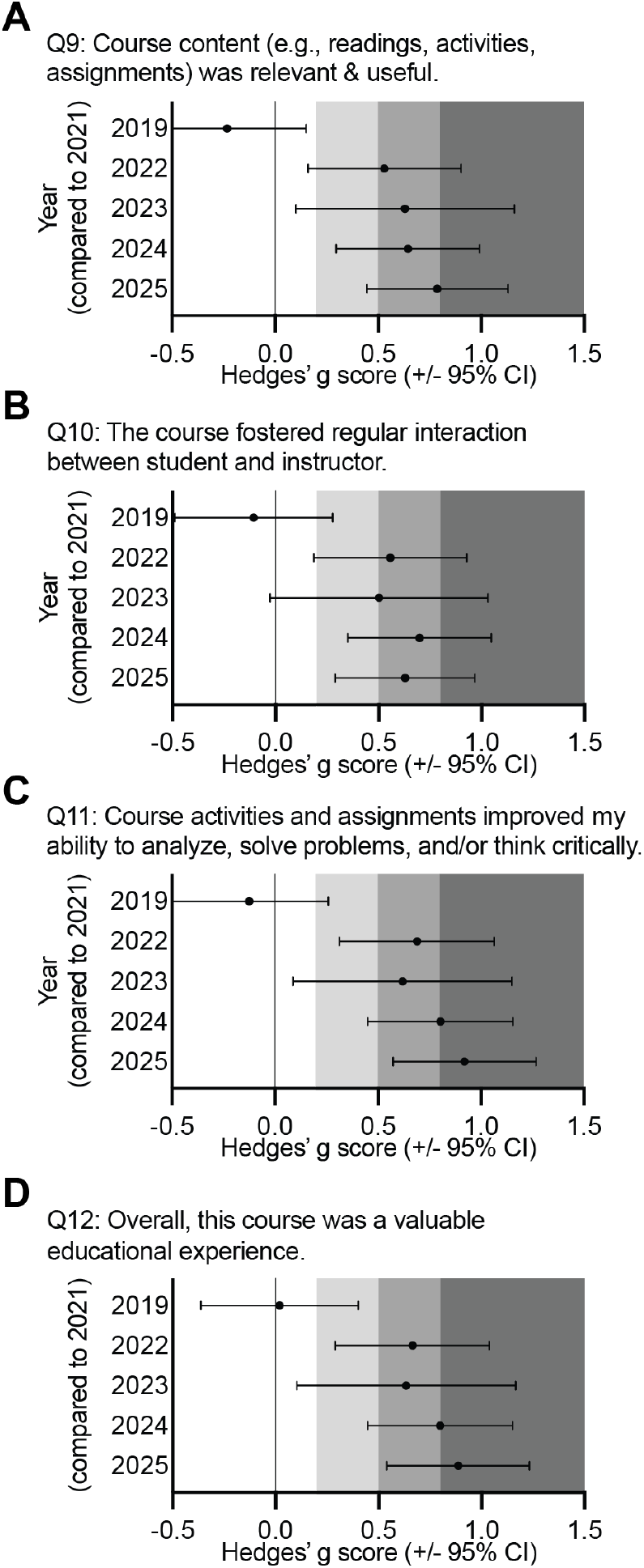
Effect size analysis of course-related evaluation outcomes. Hedges’ g effect sizes (± 95% confidence intervals) for course-related evaluation items relative to the 2021 baseline are shown for (A) Q9, (B) Q10, (C) Q11, and (D) Q12. Positive values indicate improvement relative to baseline. Comparisons between 2019 and 2021 demonstrate minimal pre-intervention differences, whereas post-intervention years (2022–2025) show moderate to large effect sizes, indicating meaningful improvements following implementation of case-based instruction.

For questions 10-12, a similar pattern was observed (Figures 3B-3D). Comparisons between 2019 and 2021 showed negligible differences, whereas comparisons between 2021 and post-intervention years demonstrated moderate to large effect sizes. Notably, for questions 11 and 12, effect sizes in 2024 and 2025 exceeded 0.8, indicating large improvements in perceived critical thinking and overall course value (Figures 3C and 3D). These findings support the conclusion that the intervention produced not only statistically significant but also educationally meaningful improvements in course-related outcomes.

### Instructor-related evaluation metrics also improved following the intervention

Instructor-related evaluation scores improved significantly following the introduction of case-based instruction. Scores for questions 3-8 were similar between 2019 and 2021, indicating consistent baseline performance prior to the intervention (Figures 4A-4F). However, comparison of 2021 to 2022 and subsequent years revealed statistically significant increases across all instructor-related measures, including enthusiasm, clarity, responsiveness, learning environment, feedback, and perceived contribution to learning. In several instances, post-intervention scores exceeded college-level averages, suggesting that the instructional changes not only improved student perceptions but did so relative to broader institutional benchmarks. The consistency of these improvements across multiple instructor-related domains indicates that the intervention had a broad impact on how students experienced instruction.

**Figure 4.**
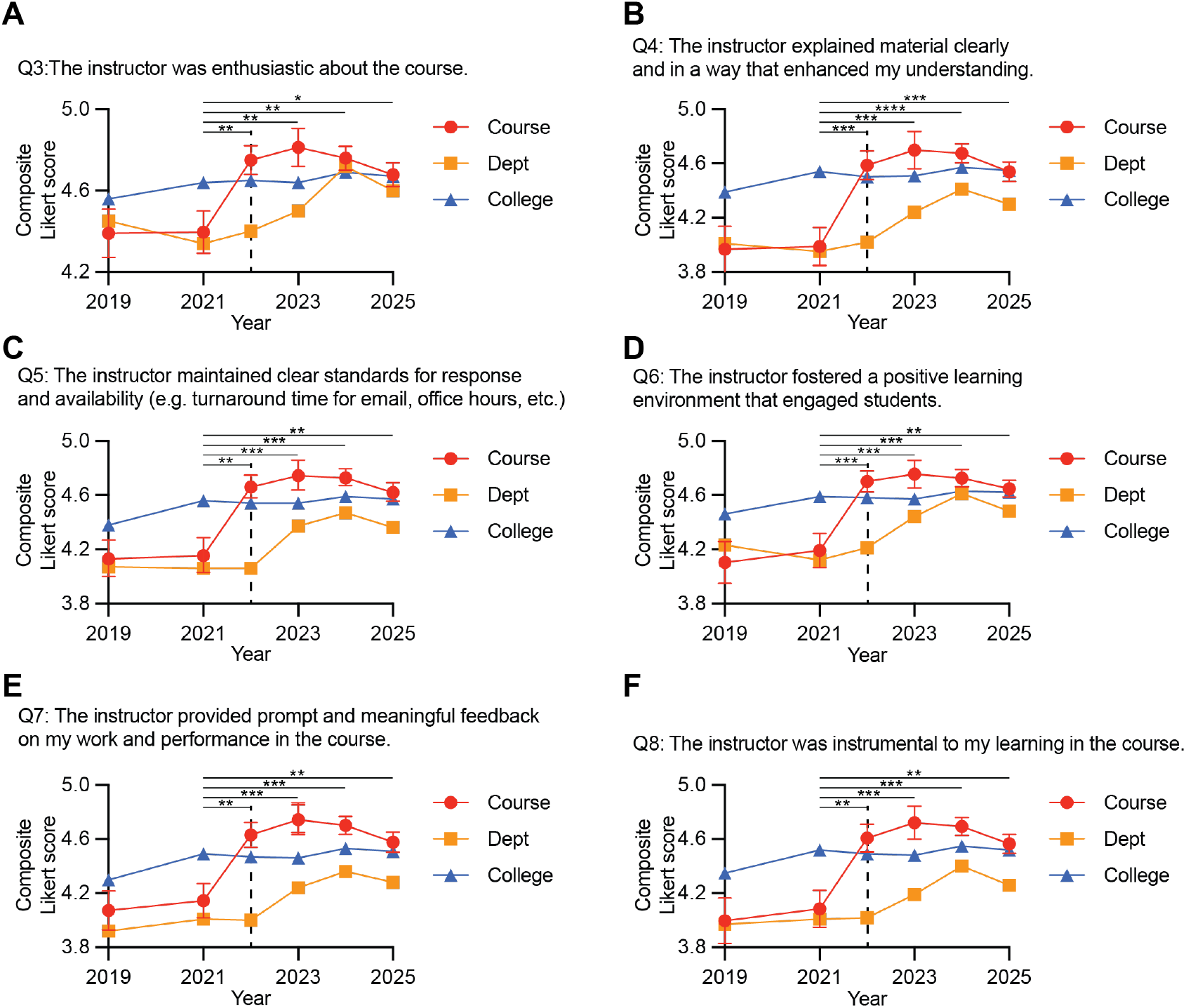
Instructor-related evaluation scores improved following implementation of case-based instruction. Composite Likert scores (mean ± SD, 1-5) for instructor-related evaluation items are shown across years for (A) enthusiasm (Q3), (B) clarity of instruction (Q4), (C) standards and availability (Q5), (D) learning environment (Q6), (E) feedback (Q7), and (F) contribution to learning (Q8). Scores represent averages across all instructors for each course offering. Course-level scores (mean and standard deviation) are compared with departmental and college averages. Case-based instruction was introduced in 2022 (dashed line). Statistical significance was determined using Welch’s t-tests comparing each year to the 2021 baseline (*p < 0.05, **p < 0.01, ***p < 0.001, ****p < 0.0001).

### Effect size analysis confirms moderate to large improvements in instructor-related outcomes

Effect size analysis further supports the impact of the intervention on instructor-related metrics. For all instructor-related questions (3-8), comparisons between 2019 and 2021 yielded effect sizes near zero, indicating no meaningful difference prior to the intervention (Figures 5A-5F). In contrast, comparisons between 2021 and post-intervention years demonstrated small to moderate effect sizes for question 3 and moderate to large effect sizes for questions 4-8. In particular, clarity of instruction (question 4) and related measures showed consistent moderate to large effect sizes across multiple years, highlighting the role of case-based teaching in enhancing student understanding. These findings indicate that the intervention meaningfully improved multiple dimensions of instructional effectiveness, reinforcing the results observed in course-level metrics.

**Figure 5.**
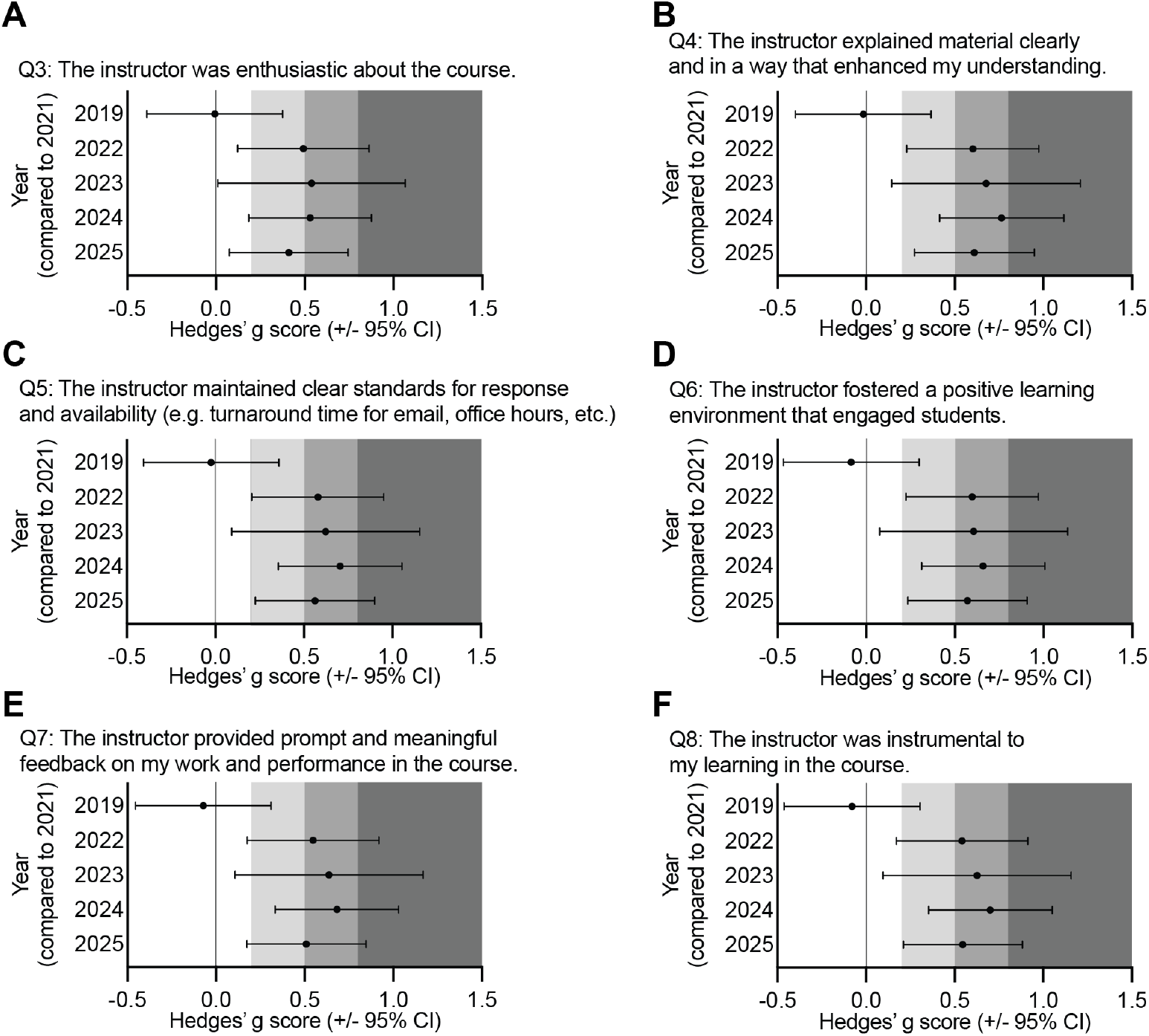
Effect size analysis of instructor-related evaluation outcomes. Hedges’ g effect sizes (± 95% confidence intervals) for instructor-related evaluation items relative to the 2021 baseline are shown for (A) Q3, (B) Q4, (C) Q5, (D) Q6, (E) Q7, and (F) Q8. Positive values indicate improvement relative to baseline. Minimal differences were observed prior to the intervention (2019 vs. 2021), whereas post-intervention years demonstrate small to large effect sizes across multiple domains, indicating consistent improvements in perceived instructional effectiveness following implementation of case-based teaching.

### Improvements in student performance paralleled evaluation outcomes and are inconsistent with grade inflation

Student performance, as measured by final course grades, improved following implementation of case-based instruction (Figure 6). Mean grades decreased from approximately 90% in 2019 to 86% in 2021 but increased to 90% in 2022 and continued to rise to 94% in 2024 before declining slightly to 92% in 2025. This pattern indicates a recovery and subsequent improvement in student performance following the instructional redesign.

**Figure 6.**
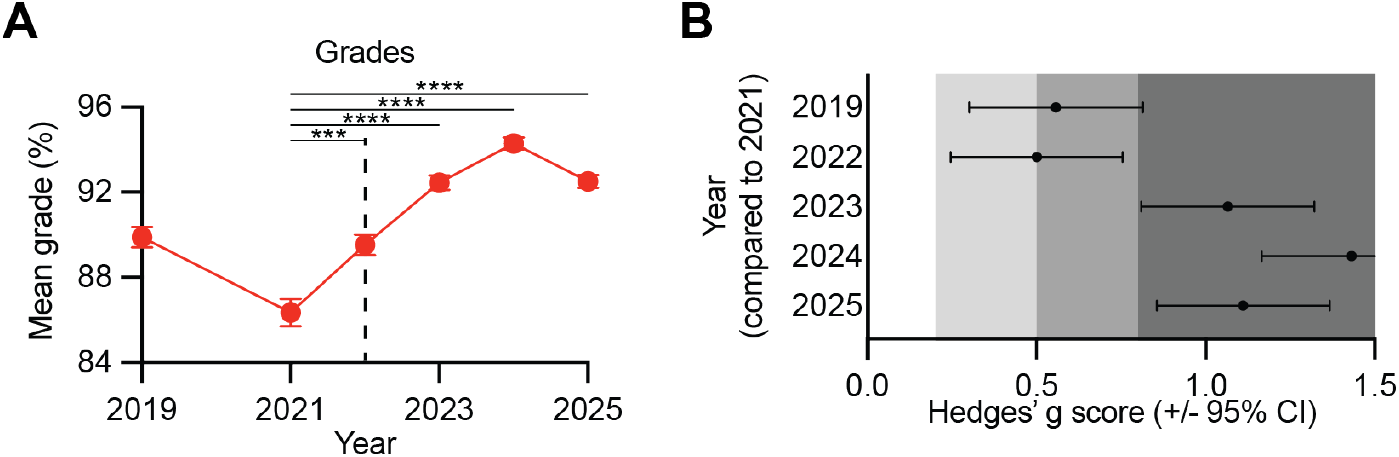
Student performance improved following implementation of case-based instruction. (A) Mean final course grades (%) across years. Points represent mean values and error bars indicate standard error of the mean (SEM). (B) Hedges’ g effect sizes (± 95% confidence intervals) for grades relative to the 2021 baseline. Assessment structure (10 quizzes and 4 examinations) and grading criteria remained consistent across all years, and quiz and examination questions were largely unchanged. Following introduction of case-based instruction in 2022, mean grades increased and remained elevated through 2025. The modest decline observed in 2025, despite sustained high evaluation scores, suggests that improvements in performance were not attributable to grade inflation.

Effect size analysis of grades showed moderate increases in 2019 and 2022 relative to 2021, and large increases in 2023–2025, with Hedges’ g values exceeding 1 in later years (Figure 6B). Importantly, the slight decline in grades observed in 2025, despite continued high evaluation scores, argues against grade inflation as the primary explanation for improved outcomes. Instead, these findings suggest that the intervention enhanced student learning in a sustained manner, resulting in improved performance without continuous upward drift in grades.

## Discussion

This study demonstrates that incorporation of clinically oriented case-based instruction into a lecture based veterinary microbiology course was associated with immediate and sustained improvements in both student perceptions and academic performance. Following implementation of the intervention in 2022, course-related and instructor-related evaluation metrics increased across nearly all measured domains and remained elevated through 2025. These gains were accompanied by improved course grades, suggesting that the instructional redesign enhanced not only student satisfaction but also measurable learning outcomes.

A notable strength of this natural experiment is that the major change across years was the method of content delivery rather than course content, learning objectives, or assessments. The 2021 offering, taught by the same coordinator and largely the same instructional team using the pre-existing format, provided an appropriate pre-intervention baseline. Similar evaluation scores in 2019 and 2021 further support the interpretation that the observed improvements were not simply part of an ongoing secular trend. Instead, the abrupt increase beginning in 2022 is temporally consistent with the introduction of case-based teaching.

These findings align with broader educational literature indicating that active and applied learning strategies can improve engagement and knowledge retention in STEM and health-professions education, and demonstrate improvement in a large lecture based setting.^7,11–13^ Traditional microbiology instruction often emphasizes memorization of organism characteristics, virulence factors, and disease associations. While foundational knowledge remains essential, learners may struggle to integrate isolated facts into clinically useful frameworks. The case examples used in this course required students to distinguish relevant from irrelevant information, interpret ambiguity, and apply microbiological principles to diagnosis and management. This likely promoted deeper cognitive processing and helped students organize content around authentic clinical problems.^8,9,18,24^ The progressive disclosure format likely contributed to these outcomes by requiring students to iteratively refine hypotheses and distinguish relevant from extraneous information, mirroring authentic clinical reasoning processes.^14^

Importantly, improvements were observed not only in course-value items such as relevance, critical thinking, and overall educational experience, but also in instructor-focused domains including clarity, enthusiasm, feedback, and contribution to learning. This suggests that students may perceive instructors more favorably when teaching methods facilitate understanding and engagement. In other words, some aspects of instructor evaluation may reflect pedagogical effectiveness as much as individual teaching style.

The concurrent rise in grades strengthens the argument that the intervention benefited learning. If higher evaluations were driven solely by lenient grading, one might expect continuous upward drift in grades over time. Instead, grades improved after the intervention, peaked in 2024, and declined modestly in 2025 while evaluations remained high. This pattern is more consistent with enhanced learning than simple grade inflation. These results contrast with Grauer et al, who did not see significant differences between lecture based and case-based/problem-based teaching within a third-year veterinary medicine course, likely due to differences in inplementation.^23^ In the Grauer study, both student cohorts had access to both lecture and case-study materials, and the lecture-based cohorts were introduced to the case-studies during lecture, but did not have problem solving/discussion sessions. Our results suggest that the introduction of case-studies within a lecture setting alone improves student outcomes, which likely minimized the difference between Grauer’s case-based and lecture-based cohorts.

This study has several limitations. It was conducted at a single institution within one veterinary curriculum, which may limit generalizability. Because this was a retrospective quasi-experimental study, unmeasured cohort differences between years cannot be excluded. Response rates varied across years, particularly in 2023, which may introduce response bias. In addition, final course grades are imperfect proxies for learning and may reflect multiple factors beyond knowledge acquisition. Despite these limitations, the consistency of findings across multiple years and across numerous independent evaluation domains supports the robustness of the observed effect. Future studies could prospectively compare specific case designs, measure knowledge retention longitudinally, or assess downstream clinical reasoning performance during later veterinary training. Additionally, this while data did not investigate the effect of case-studies across different student demographics, we expect the clinical case studies improve student course perception in part by strengthening science identity^25^.

In conclusion, integrating clinically rich, distractor-containing case examples into a content-dense veterinary microbiology course was associated with sustained improvements in student engagement, perceptions of instructional quality, and academic performance. These findings support case-based teaching as a practical and scalable strategy for improving microbiology education in veterinary and other health-professions curricula.

## Acknowledgements

The authors thank Chloe Van Horn for technical assistance with data organization. We also thank Christopher Stanton, Kia Hendrix, and John Michael Jordi for their assistance in providing access to data used in this study. This study was supported by a grant from the National Institutes of Health (1R21AI204394 to A.E.), University of Florida Office of Research and College of Veterinary Medicine.

---

**Supplemental figure S1.**
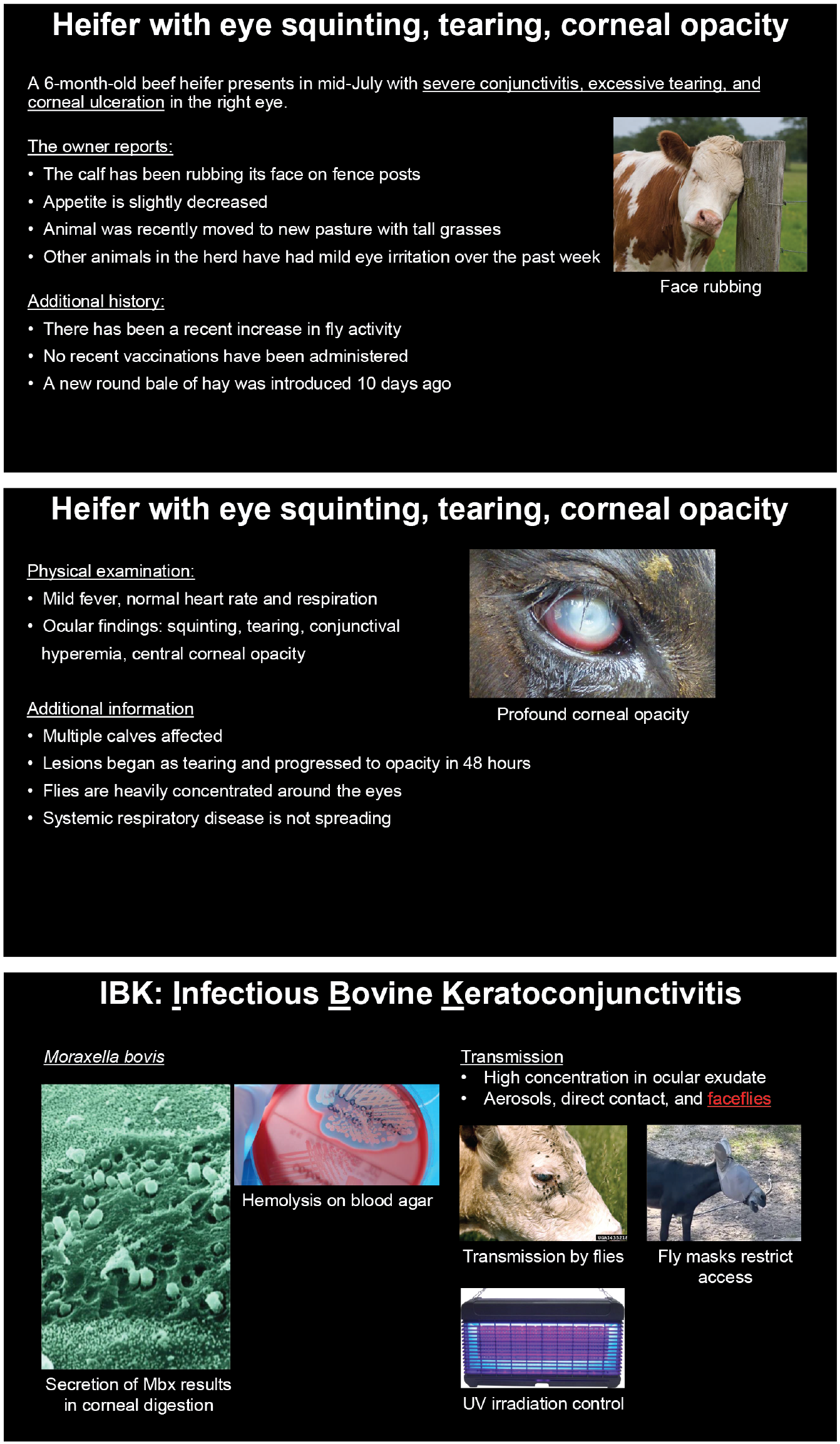
Example of progressive clinical case-based instruction used in veterinary microbiology lectures. Representative slides illustrating the stepwise presentation of a clinical case of infectious bovine keratoconjunctivitis caused by *Moraxella bovis*. Initial slides provide clinical history and environmental context, including both relevant and extraneous information, requiring students to differentiate potential infectious and non-infectious causes. Subsequent slides introduce physical examination findings and epidemiological clues, including involvement of multiple animals and progression of corneal lesions, guiding students toward an infectious etiology. The final slide reveals the causative organism and integrates key microbiological concepts, including virulence mechanisms (e.g., RTX toxin-mediated corneal damage), diagnostic features, transmission pathways, and control strategies. This progressive disclosure format is used to promote active learning, clinical reasoning, and application of microbiological principles.

## Notes

### Competing Interest Statement

The authors have declared no competing interest.

